# Modulation of *Salmonella* virulence by a novel SPI-2 injectisome effector that interacts with the dystrophin-associated protein complex

**DOI:** 10.1101/2023.12.07.570645

**Authors:** Xiu-Jun Yu, Haixia Xie, Yan Li, Mei Liu, Ruhong Hou, Alexander V. Predeus, Blanca M. Perez Sepulveda, Jay C. D. Hinton, David W. Holden, Teresa L. M. Thurston

## Abstract

The injectisome encoded by *Salmonella* pathogenicity island 2 (SPI-2) had been thought to translocate 28 effectors. Here, we used a proteomic approach to characterise the secretome of a clinical strain of invasive non-typhoidal *Salmonella enterica* serovar Enteritidis, that had been mutated to cause hyper-secretion of the SPI-2 injectisome effectors. Along with many known effectors, we discovered the novel SseM protein. *sseM* is widely distributed between the five subspecies of *Salmonella enterica,* is found in many clinically-relevant serovars, and is co-transcribed with *pipB2, a* SPI-2 effector gene. Translocation of SseM required a functional SPI-2 injectisome. Following expression in human cells, SseM interacted with five components of the dystrophin-associated protein complex (DAPC), namely β-2-syntrophin, utrophin/ dystrophin, α-catulin, α-dystrobrevin and β-dystrobrevin. The interaction between SseM and β-2-syntrophin and α-dystrobrevin was verified in *S.* Typhimurium-infected cells and relied on the PDZ domain of β-2-syntrophin and a sequence corresponding to a PDZ-binding motif (PBM) in SseM. A Δ*sseM* mutant strain had a small competitive advantage over the wild-type strain in the *S.* Typhimurium/mouse model of systemic disease. This phenotype was complemented by a plasmid expressing wild type SseM from *S.* Typhimurium or *S.* Enteritidis and was dependent on the PBM of SseM. Therefore, a PBM within a *Salmonella* effector mediates interactions with the DAPC and modulates systemic growth of bacteria in mice.

**Importance:** In *Salmonella enterica*, the injectisome machinery encoded by *Salmonella* pathogenicity island 2 (SPI-2) is conserved among the five subspecies and delivers proteins (effectors) into host cells that are required for *Salmonella* virulence. The identification and functional characterisation of SPI-2 injectisome effectors advances our understanding of the interplay between *Salmonella* and its host(s). Using an optimised method for preparing secreted proteins and a clinical isolate of the invasive non-typhoidal (iNTS) *Salmonella enterica* serovar Enteritidis strain D24359, we identified 22 known SPI-2 injectisome effectors and one new effector - SseM. SseM modulates bacterial growth during murine infection and has a sequence corresponding to a PDZ-binding motif that is essential for interaction with the PDZ-containing host protein β-2-syntrophin and other components of the dystrophin-associated protein complex (DAPC). To our knowledge, SseM is unique among *Salmonella* effectors in containing a functional PDZ-binding motif and is the first bacterial protein to target the DAPC.

## Introduction

Following entry into host cells, *Salmonella enterica* serovar Typhimurium (*S.* Typhimurium) resides in membrane-bound compartments known as *Salmonella-*containing vacuoles (SCVs). Acidification and nutritional starvation of the vacuole lumen activate the two-component regulatory system SsrAB to induce expression of *Salmonella* pathogenicity island 2 (SPI-2) genes followed by the assembly of a type three secretion apparatus known as the SPI-2 injectisome [1–3]. The associated gatekeeper complex, comprising SsaL, SsaM and SpiC, enables the injectisome to secrete the translocon proteins SseBCD while preventing the premature translocation of effectors; once the translocon pore is formed on the SCV membrane, the neutral pH of host cell cytosol is sensed and the signal is transduced into the bacterial cytosol to disassociate the gatekeeper complex from the export gate component SsaV to allow translocation of effectors [4–6]. Approximately 28 such effectors are translocated into host cells [7, 8]; collectively these enable bacterial replication in host cells and suppress both innate and adaptive immune responses [1, 9-11].

Since the discovery of SPI-2 injectisome [9, 10, 12], different approaches have been explored for the identification of SPI-2 injectisome effectors. By similarity search for known effectors of other injectisomes, SspH1/2 [13]; SlrP, SifA, SseI, SseJ and SifB [14, 15]; SopD2 [16], PipB2 [17], and SseK1/2 [18] were shown to be SPI-2 injectisome effectors. Screening for SsrAB-regulated factors (*srf*) either by MudJ mutagenesis [19] or transcriptomic analysis [20, 21] revealed SrfA-M and SseL. Both SrfH (i.e. SseI) and SseL were subsequently verified as SPI-2 effectors. Screening a transposon mini-Tn5-cycler-generated library of translational fusions between *Salmonella* chromosomal genes and *cyaA*’ during cell infection, identified known effectors (SlrP, PipB2, SseJ, SrfH and AvrA) and new effectors (SteA, SteB and SteC) [22]. A gatekeeper mutant Δ*ssaL* strain that hypersecretes effectors into culture medium has been used to identify effectors by proteomic analysis: 17 known effectors and 6 new SPI-2 effectors were identified: SpvD, GtgE, GtgA, SteD, SteE and CigR [23]. Both the CyaA translocation screen and the Δ*ssaL*-based secretion screen used *S.* Typhimurium strain 12023 (i.e. ATCC14028) and its derivatives [22] [23]. In an alternative approach, Auweter et al [24] used stable isotope labelling with amino acids in cultures of *S.* Typhimurium SL1344 wt and SPI-2 null mutant strains to identify 12 effectors: SpvC/D, SopD2, SifA, SseJ, SteC, SteA, SseL, PipB2, PipB, GtgE and SteE.

*S.* Typhimurium strain ATCC14028 and SL1344 have been widely used in laboratories for over 50 years; the former was isolated from a chick in 1960 and the latter from cattle in 1966. Proteomic analysis has not yet been used to screen SPI-2 effectors from contemporary clinical isolates. In this study, we sought to identify SPI-2 effectors from a clinical isolate of invasive non-typhoidal *Salmonella* (iNTS) Enteritidis strain D24359 by comparing the secretomes of an isogenic hypersecretion gatekeeper mutant and a SPI-2 null mutant. We found 22 known effectors and one previously unidentified effector - SseM. SseM is present in all 5 subspecies of *S. enterica*. The postsynaptic density-95/discs large/zonula occludens-1 (PDZ) domain binding motif (PBM) of SseM mediates interaction with dystrophin-associated protein complex (DAPC) and is involved in modulation of *Salmonella* virulence.

## Results

### Discovery of SPI-2 injectisome effector SseM by proteomic analysis

To investigate the SPI-2 injectisome effector repertoire of a clinical isolate of *Salmonella enterica* subspecies *enterica,* we exploited the hypersecretion phenotype of a gatekeeper mutant. We chose an invasive non-typhoidal *Salmonella enterica* serovar Enteritidis (*S.* Enteritidis) strain D24359 that was isolated from blood of a Malawian patient and is sensitive to antibiotics carbenicillin, kanamycin and chloramphenicol [25] to make a Δ*spiC* single mutant (an effector hypersecretion mutant) and a Δ*spiC,ssaC* SPI-2 null mutant [4]. The bacterial strains were grown in SPI-2-inducing medium MgM-MES at pH 5.0 for 6 h. Then the supernatant was concentrated and subjected immediately to SDS-PAGE after which a 1 cm gel slice was analysed by mass spectrometry (**Fig. 1A**). The resulting peptides were compared to predicted protein sequences from the annotated *S.* Enteritidis strain D24359 sequence. From two experiments, we identified 22 known SPI-2 effectors and one previously unidentified effector D24359_01053 (**Table 1 and Fig. 1B**).

**Fig.1.**
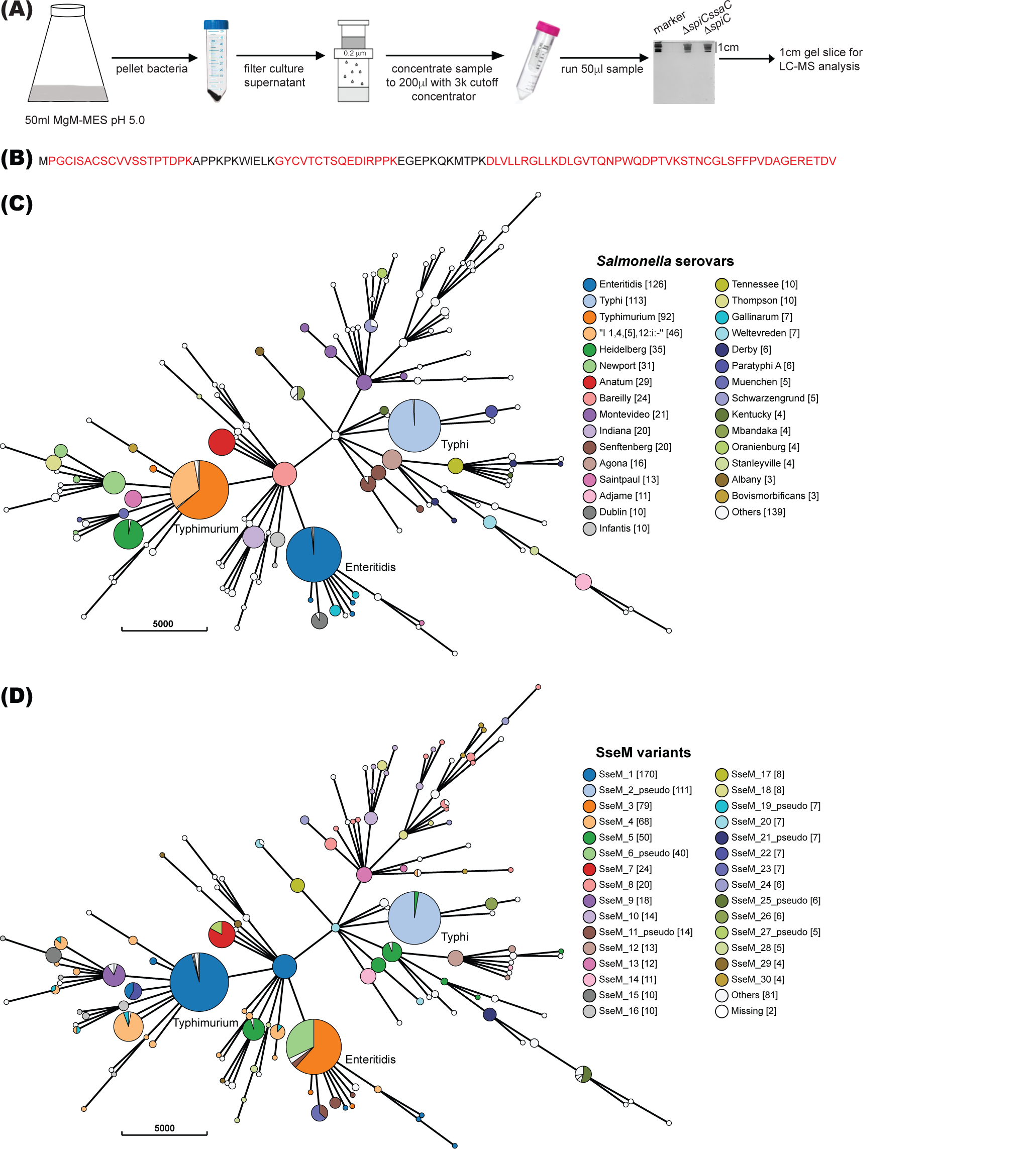
Identification of SPI-2 effector SseM and conservation in *Salmonella enterica* subspecies I genomes. (A) Schematic for identification of novel *Salmonella* secreted proteins (B) Matched peptides (red fonts) of newly identified effector, D24359_01053, from one MS analysis. (C-D) GrapeTree phylogenetic visualization of SseM distribution and protein sequence variation; branch length indicates the number of allele differences between the cgMLST types, as shown by the scale bar. The nodes with fewer than 900 allele differences were collapsed into bubbles, which are consistent with serovars. The size of each bubble is proportional to the number of genomes it represents. The bubbles that correspond to *Salmonella* serovars Typhi, Typhimurium and Enteritidis are labelled. The colour of the bubbles indicates the serovars (C) or the diverse SseM protein sequence types (D).

**Table 1.**
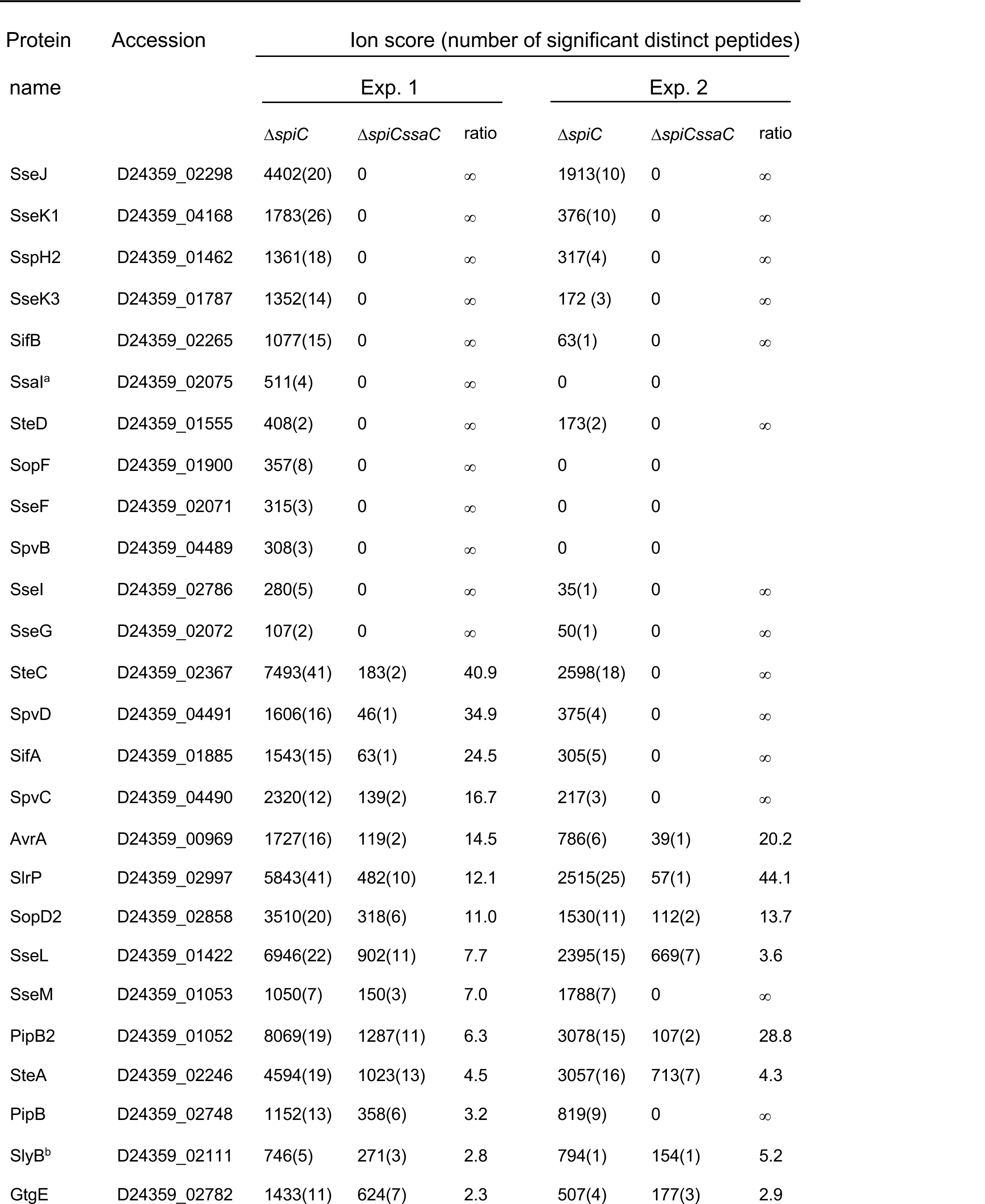
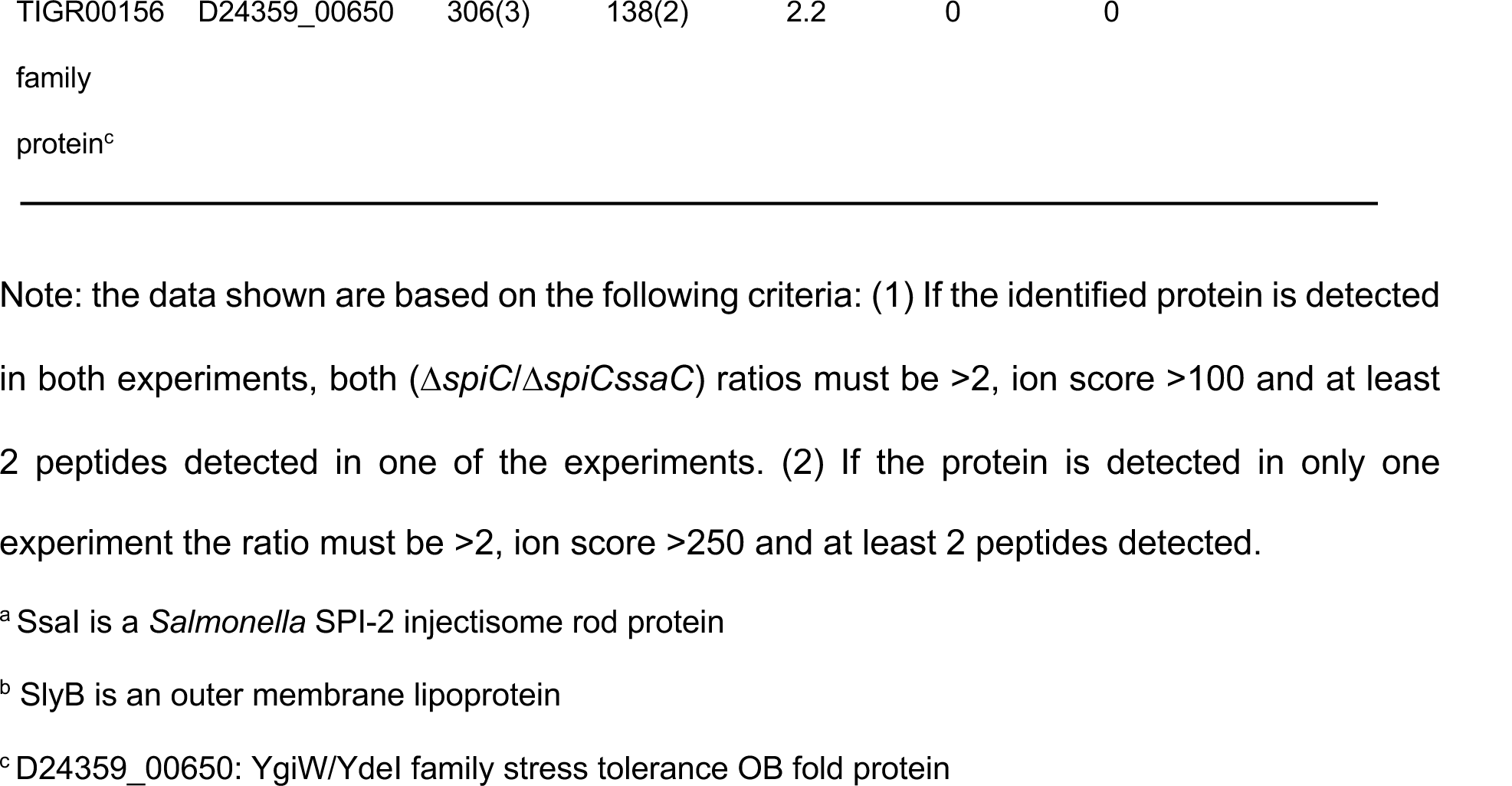
Identified secreted proteins with a ratio of ion scores (Δ*spiC*/Δ*spiCssaC*) >2.

D24359_01053 consists of 103 amino acids. Searching the BLAST Protein database of *S.* Typhimurium LT2 showed that D24359_01053 is almost identical to STM2779: there are 95 identical residues in the predicted 110 amino acids of STM2779 (**sFig. 1A**). Bioinformatic analysis revealed that D24359_01053/STM2779 is conserved in all five subspecies of *Salmonella enterica* but not in *S. bongori*, with most being predicted to be 110 amino acids in length (**sFig. 1B**). We named this effector SseM (*<u>S</u>almonella* <u>s</u>ecreted <u>e</u>ffector M).

To further analyse the distribution of SseM and SseM variants among serovars of *S*. *enterica* subspecies *enterica*, 834 complete *Salmonella* genomes with the highest assembly quality were downloaded from Enterobase (https://enterobase.warwick.ac.uk/) and used to construct an *sseM* database with *stm2779* as the reference sequence. The SseM protein variants in the context of genomic or serovar diversity were displayed with the Grapetree phylogenetic tool [26] (**Fig. 1C and D**). Most of the sequenced strains of different serovars had full length SseM (**Fig. 1D**), with the N- and C-terminal regions of SseM highly conserved (**sFig. 2**). Our analysis revealed the presence of two common pseudogene variants; *sseM*_2_pseudo, present in 111 out of 113 *S.* Typhi genomes and *sseM*_6_pseudo, present in 40 out of 126 *S.* Enteritidis genomes (**Fig. 1D**). *sseM*_2_pseudo is the result of an additional cytosine in *sseM* of *S*. Typhi, which generates a stop codon immediately after the predicted 30^th^ residue; while the *sseM*_6_pseudo is due to the mutation of the predicted 84^th^ codon TGG to TAG, resulting in a truncated version of SseM (**sFig. 3**). In summary, SseM is widely distributed among the five subspecies of *Salmonella enterica* and so represents a new conserved “core” effector protein.

### Expression, secretion, and translocation of SseM

*sseM* is located 175 nt downstream of SPI-2 effector gene *pipB2* (**Fig. 2A**). Both *pipB2* and *sseM* (*stm2779*) share the same transcriptional start site [27] and are controlled by SsrAB [20, 28]. To verify SsrAB dependence on expression of SseM and to test if *pipB2* and *sseM* were operonic or not, a rabbit polyclonal antibody against the C-terminal peptide (PYFPVVPGERETDV) of *S.* Typhimurium SseM was obtained. *S.* Typhimurium 12023 wt and derivative strains were grown in MgM-MES at pH 5.0, and proteins in whole bacterial lysates were immunoblotted. The antibody detected SseM in wt and a Δ*sseM* mutant expressing SseM from a plasmid (p*sseM*) but not in lysates derived from the Δ*sseM* mutant, demonstrating the specificity of the SseM antibody (**Fig. 2B**). As expected, SseM was not detected in lysates derived from a *ssrA::mTn5* mutant. Furthermore, deleting the promoter of *pipB2* but not *pipB2* itself led to the loss of SseM (**Fig. 2B**). Taken together, the data indicate that *sseM* and *pipB2* are bicistronic, with their expression activated by SsrAB.

**Fig.2.**
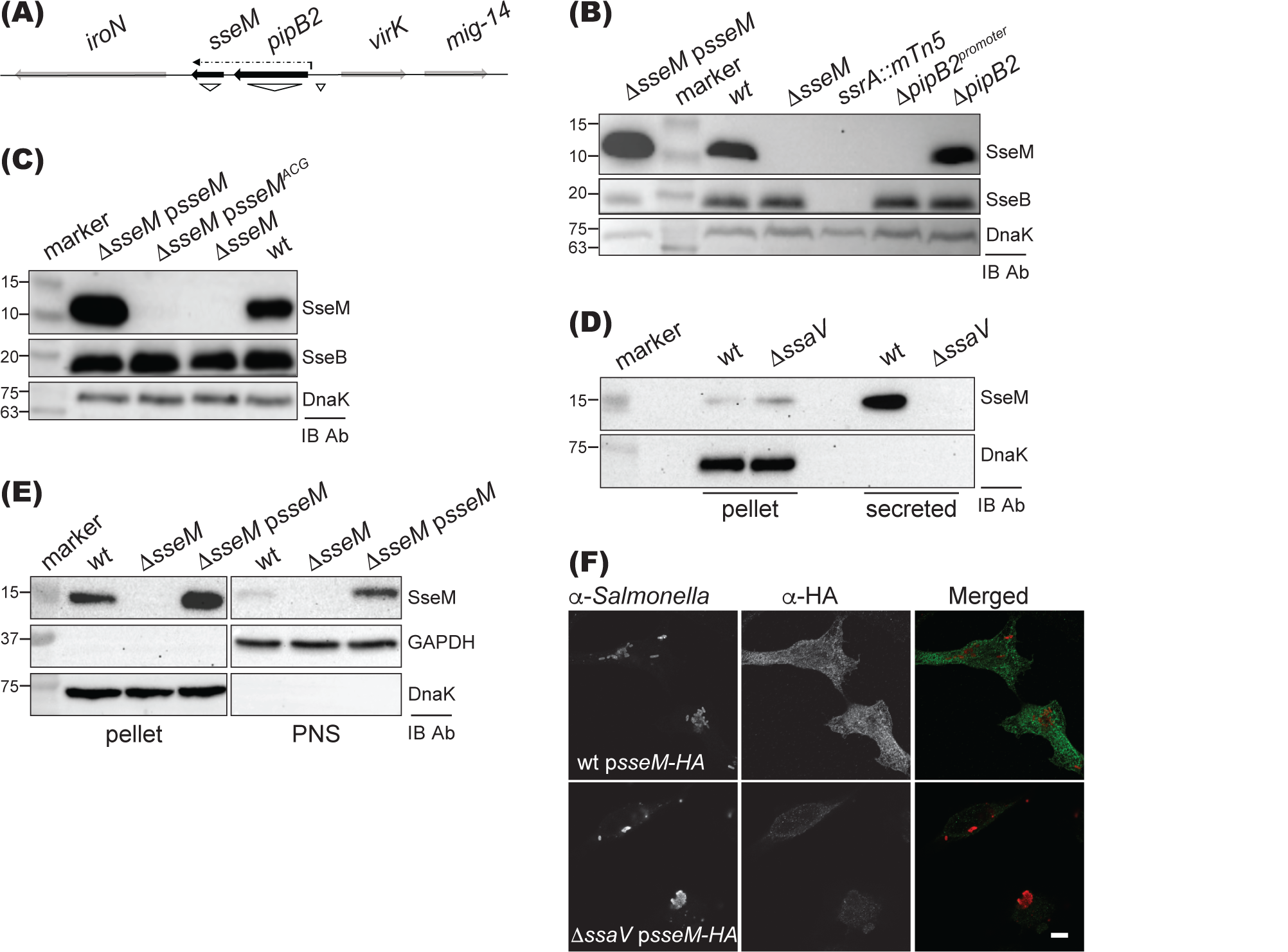
Analysis of expression, secretion and translocation of SseM. (A) Genetic organisation of *pipB2*-*sseM* operon and regions of deletion used in (B) are indicated with ∇. The dash dotted arrow indicates the transcript of *pipB2-sseM*. (B) Expression of SseM is controlled by the *pipB2* promoter and SsrAB. Bacterial strains were grown in MgM-MES at pH 5.0 for 6h. Bacterial lysates were analysed by immunoblotting. Intrabacterial protein DnaK and SPI-2 translocon protein SseB were used as controls. (C) Indicated bacterial lysates, including strains expressing the mutation of the predicated start codon ATG to ACG (p*sseM^ACG^*) were analysed by immunoblotting. (D) Secretion of SseM upon pH shift. Bacterial strains were grown at pH 5.0 for 4 h, then switched pH to 7.2 for another 1.5 h before preparing samples for immunoblotting. (E) Translocation analysis by immunoblotting. HeLa cells were infected for 5 h, then proteasome inhibitor MG132 was added for another 3h before fractionation with Triton-X 100. Triton-X 100 soluble fraction (PNS) contains translocated proteins. (F) Translocation analysis by confocal microscopy. HeLa cells were infected with bacteria expressing SseM-HA for 5 h, then proteasome inhibitor MG132 was added for another 3 h before fixation. Fixed cells were labelled with antibodies to visualize *Salmonella* (red) and SseM-HA (green). Scale bar: 5 μm.

As SseM from *S.* Typhimurium strain LT2 and most of *S. enterica* species have been annotated as a 110-residue long protein that uses TTG rather than ATG (21 nucleotides downstream of TTG), as its start codon (**sFig. 4**), we defined the actual start codon of *sseM* by changing the ATG to ACG on plasmid p*sseM*. The resulting plasmid p*sseM^ACG^* was transformed into the Δ*sseM* mutant strain to check the expression of SseM by immunoblot. SseM was undetectable in the Δ*sseM* mutant carrying plasmid p*sseM^ACG^*, indicating that *sseM* uses the ATG as its start codon to encode a 103-residue long protein (**Fig. 2C**).

Next, we investigated SPI-2 injectisome-dependent secretion of SseM by immunoblotting. The wt and a Δ*ssaV* mutant strains were grown in MgM-MES at pH 5.0 for 4 h to assemble the SPI-2 injectisome, then the pH of medium was changed to 7.2 to allow effector secretion [5]. Although SseM was detected in bacterial lysates from both wt and Δ*ssaV* mutant strains, secreted SseM was only detected in samples prepared from the wt culture (**Fig. 2D**). This result agrees with the mass spectrometric result of *S.* Enteritidis strain D24359, demonstrating that SseM secretion is dependent on the SPI-2 injectisome.

Translocation of SseM into mammalian cells from intracellular *Salmonella* was then tested by immunoblotting. For this, HeLa cells were infected with different bacterial strains for 8 h, then translocated proteins were extracted from post-nuclear supernatant with Triton X-100 and subject to immunoblotting using the anti-SseM antibody. A small quantity of translocated SseM was detected from HeLa cells infected with wt strain, and significantly more was detected in the mutant strain carrying *sseM* on a plasmid (**Fig. 2E**). However, attempts to detect translocated SseM with the rabbit anti-SseM antibody by immunofluorescence microscopy failed. To further investigate translocation of SseM by microscopy, a plasmid expressing C-terminal HA-tagged SseM (p*sseM-HA*) was transformed into the wt or Δ*ssaV* mutant strains and these were used to infect HeLa cells. Translocated SseM-HA was detected with an anti-HA epitope antibody in cells infected by wt but not the Δ*ssaV* mutant strain (**Fig. 2F**). These results demonstrate that SseM is translocated into host cell via the SPI-2 injectisome.

### SseM interacts with β-2-syntrophin and its associated proteins

To identify host cell proteins with which SseM interacts, stable HeLa cell lines expressing GFP or GFP::SseM were established, then lysates were subjected to GFP-trap immunoprecipitation followed by mass spectrometric analysis. Utrophin/dystrophin, α-catulin, α-dystrobrevin, β-dystrobrevin and four PDZ domain-containing proteins β-2-syntrophin, disks large homolog 1 (DLG1), peripheral plasma membrane protein CASK and protein lin-7 homolog C were co-purified with GFP::SseM but not by GFP alone (**Table 2**). These putative targets of GFP::SseM were also specifically co-immunoprecipitated with Flag::SseM but not by another Flag-tagged SPI-2 injectisome effector (Flag::SpvD) from transiently transfected HEK 293 cells (**Table 2**). β-2-syntrophin, utrophin/dystrophin, α-catulin, α-dystrobrevin and β-dystrobrevin are components of the dystrophin-associated protein complex (DAPC) signalosome [29, 30], and had much higher ion scores and number of peptides detected in our screenings than the other three PDZ domain-containing proteins. Therefore, further work was focussed on the interaction between SseM and β-2-syntrophin and its associated proteins.

**Table 2.**
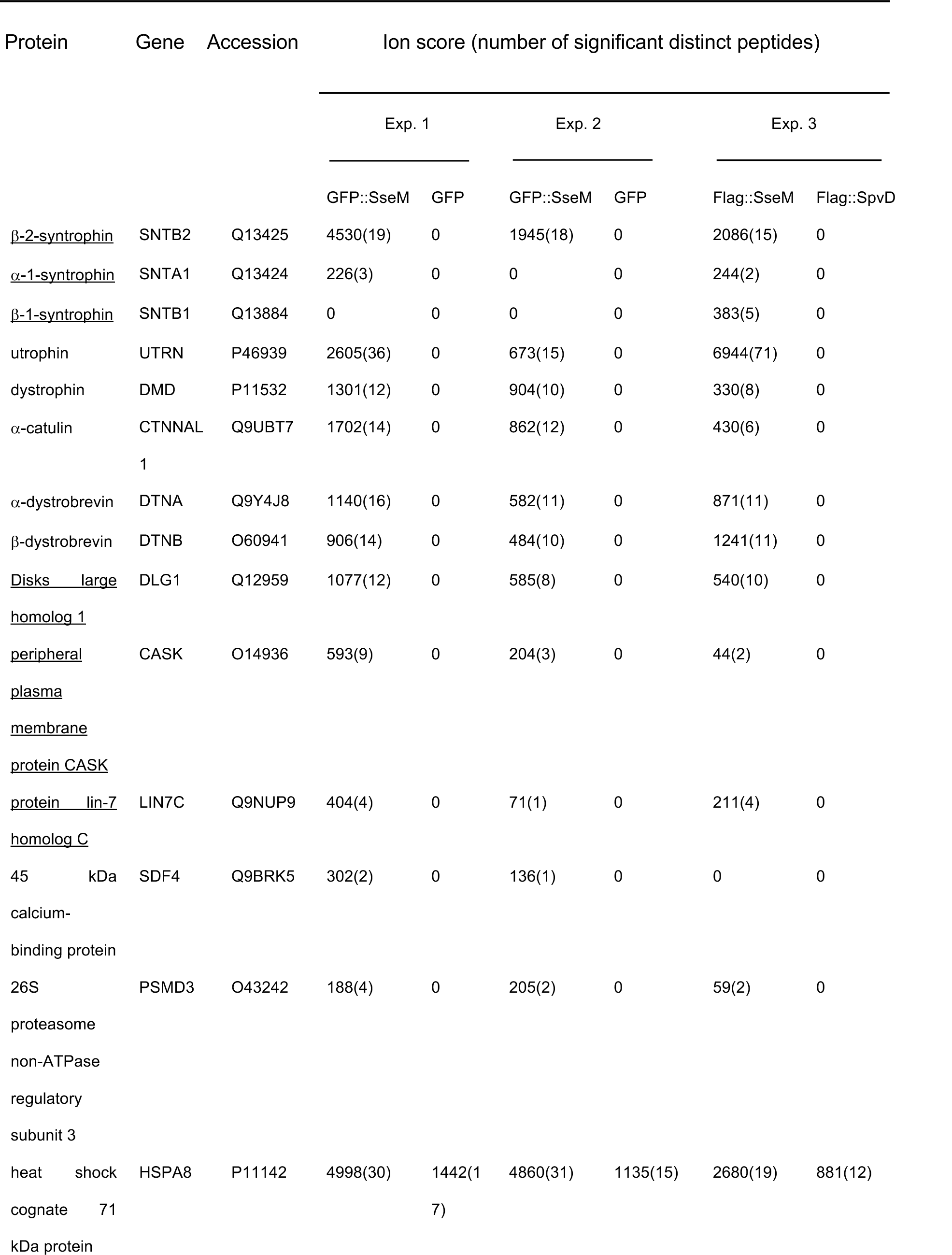
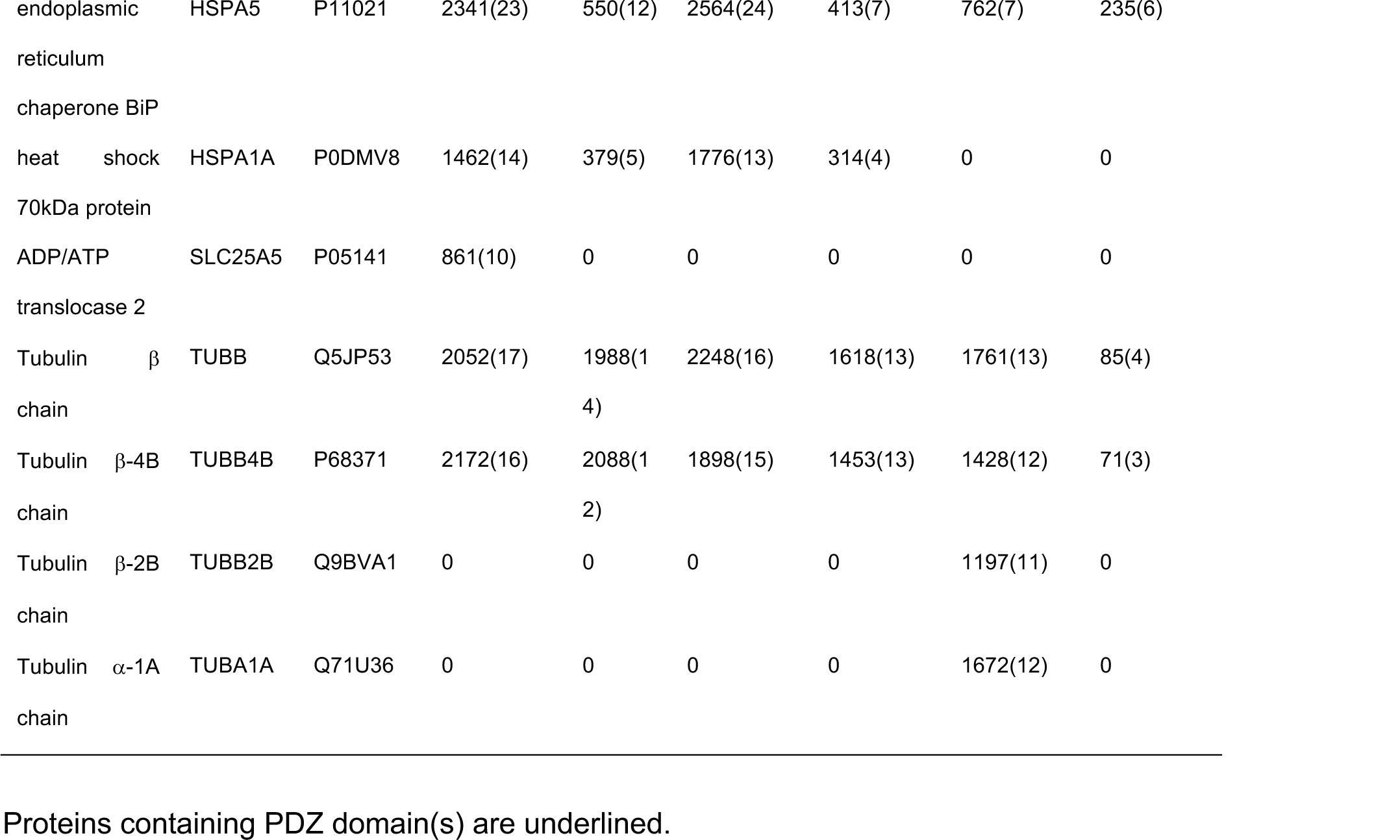
LC-MS/MS analysis of SseM interacting proteins.

As an independent test of the validity of the mass spectrometry results, and to check if SseM of *S.* Enteritidis (SseM^SEN^ to distinguish it from SseM of *S.* Typhimurium) also interacts with the same targets, HEK 293 cells were transiently transfected to express GFP-tagged effectors and cell lysates subjected to immunoprecipitation before immunoblotting. Like GFP::SseM, GFP:: SseM^SEN^ also interacted with β-2-syntrophin and α-dystrobrevin (**Fig. 3A**). However, GFP::SseM-HA failed to interact with β-2-syntrophin and α-dystrobrevin, indicating that the C-terminus of SseM is crucial for its interaction with the host cell targets.

**Fig.3.**
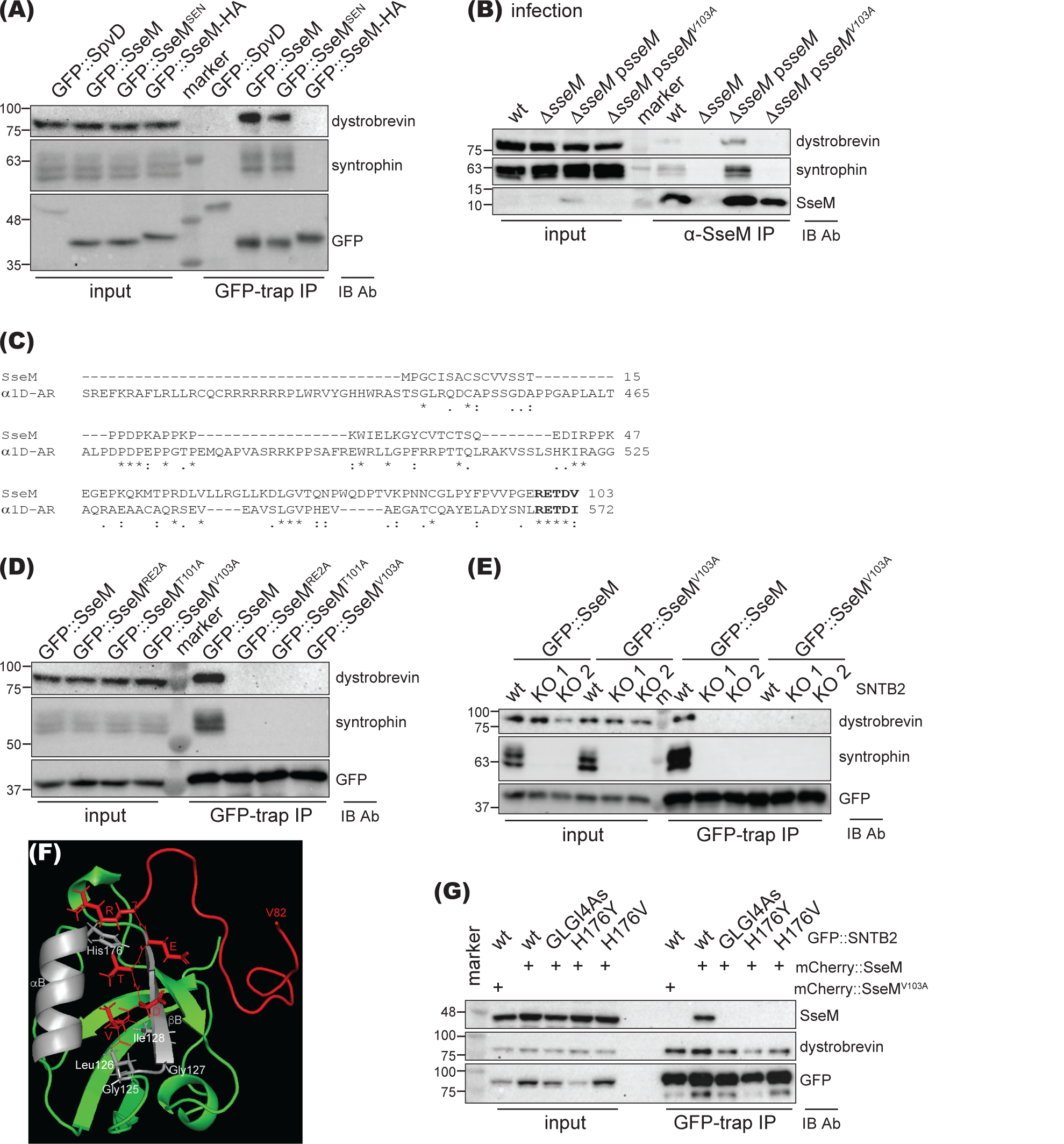
SseM interacts with β-2-syntrophin and α-dystrobrevin. (A) GFP-tagged protein was expressed in HEK293 cells, immunoprecipitated with GFP-trap agarose and analysed by immunoblotting. GFP::SpvD was used as negative control. (B) HeLa cells were infected with *Salmonella* for 14.5 h, then proteasome inhibitor MG132 was added for another 3 h. Cell lysates were immunoprecipitated with rabbit anti-SseM antibody before immunoblotting. (C) Alignment of SseM and C-terminus of α_1D_-AR. Bold fonts indicate the conserved PDZ binding motif (PBM). (D) Ectopically expressed GFP-tagged SseM variants in HEK293 cells were immunoprecipitated with GFP-trap agarose and analysed by immunoblotting. (E) WT or β-2-syntrophin (SNTB2) knockout HEK293 cells (KO1 and KO2) were transfected with GFP::SseM and protein-protein interactions analysed in cell lysates by immuno-precipitation and immunoblotting. GFP::SseM^V103A^ was included as an additional control. “m” indicates protein marker. (F) Model of complex between SseM PBM (red) and PDZ domain of β-2-syntrophin (green and grey) derived from AlphaFold Colab Multimer. α-helix B (αB) and β-strand B (βB) of β-2-syntrophin PDZ domain are coloured in grey with key amino acids annotated. (G) PDZ domain of β-2-syntrophin is required to interact with SseM. mCherry::SseM was co-expressed with indicated GFP-tagged β-2-syntrophin (GFP::SNTB2) variant in SNTB2 KO1 HEK293 cells, and subject to protein-protein interaction analysis. mCherry::SseM^V103A^ was used as control.

Then, to test if SseM translocated from intracellular *Salmonella* interacts with the same host cell proteins, HeLa cells were infected with bacterial strains for 17.5 h. Infected cells were then lysed, the lysate proteins were immunoprecipitated with the rabbit anti-SseM antibody and subjected to immunoblotting. β-2-syntrophin and α-dystrobrevin were co-immunoprecipitated from cells infected with wt strain or the Δ*sseM* mutant complemented with plasmid p*sseM* but not the Δ*sseM* mutant strain (**Fig. 3B**). Therefore, and importantly, translocated SseM interacts with β-2-syntrophin and α-dystrobrevin in physiological conditions.

### A PDZ domain-binding motif of SseM and PDZ domain of β-2-syntrophin are essential for interaction between SseM and β-2-syntrophin

Syntrophins use their PDZ domains to interact with C-terminal PDZ domain-binding motifs (PBMs) in other proteins such as the α_1D_-adrenergic receptor (α_1D_-AR) [29–31]. As SseM-HA failed to interact with β-2-syntrophin and α-dystrobrevin, we hypothesised that SseM itself might have a C-terminal PBM. Alignment of α_1D_-AR and SseM revealed that SseM indeed has a RETDV sequence at its C-terminus that corresponds to the PBM (^568^RETDI^572^) in α_1D_-AR (**Fig. 3C**). To test if the RETDV motif of SseM was required for its interaction with the host cell targets, a set of mutated GFP-SseM variants transiently expressed in HEK 293 cells were immunoprecipitated with GFP-trap and subjected to immunoblotting. As shown in **Fig. 3D**, GFP::SseM^RE2A^, GFP::SseM^T101A^, and GFP::SseM^V103A^ failed to interact with β-2-syntrophin and α-dystrobrevin. Consistent with this result, translocated SseM^V103A^ also failed to interact with β-2-syntrophin and α-dystrobrevin (**Fig. 3B**), demonstrating that the putative PBM of SseM is required for its interaction with β-2-syntrophin and α-dystrobrevin in transfected or infected cells.

To test if β-2-syntrophin mediates interaction between SseM and α-dystrobrevin, we knocked out β-2-syntrophin in HEK293 cells with two different guideRNAs (g361 and g363). Knockout of β-2-syntrophin abolished α-dystrobrevin co-immunoprecipitated with GFP-SseM (**Fig. 3E**). This result suggests that SseM interacts with its host cell targets through the interaction between its PBM and the PDZ domain of β-2-syntrophin. In agreement with this hypothesis, predication of interaction between SseM and PDZ domain of β-2-syntrophin with AlphaFold Colab Multimer showed that the RETDV residues of SseM fit in the binding pocket between β-strand B (βB) and α-helix B (αB) of the β-2-syntrophin PDZ domain (**Fig. 3F**). Based on our structural predication and the structural data of other PDZ domains and PBMs [32, 33], we predicted that residues ^125^GLGI^128^ or H176 of β-2-syntrophin (Highlighted in **sFig. 5**) are crucial for mediating its interaction with the PBM of SseM. To test this, GFP-tagged β-2-syntrophin or its variants were co-expressed with mCherry-tagged SseM or SseM^V103A^ in β-2-syntrophin knock out HEK293 cells, and cell lysates were subject to GFP-trap immunoprecipitation. Mutating GLGI to 4 As or mutating the substrate-specific residue H to Y or V of β-2-syntrophin abolished its interaction with SseM although the mutants still interacted with α-dystrobrevin (**Fig. 3G**). Taken together, the data demonstrate that the PBM of SseM and PDZ domain of β-2-syntrophin are essential for the interaction between SseM and β-2-syntrophin.

### The PBM of SseM modulates Salmonella growth during murine infection

We next assessed the contribution of SseM to *Salmonella* growth in systemic tissues of mice by competitive index (CI) analysis [34], involving intraperitoneal injection of a mixed inoculum of wt and Δ*sseM* mutant strains in susceptible mice. At 3 days post inoculation, the Δ*sseM* mutant strain significantly outcompeted the *wt::Km* strain (CI=1.800 ± 0.558). The Δ*sseM* mutant strain harbouring plasmid p*sseM* failed to outcompete the *wt::Km* strain (CI= 0.842 ± 0.196) and the CI results were significantly different to that of the Δ*sseM* mutant strain *vs wt::Km* strain (**Fig. 4**), showing that the small fitness difference was SseM-dependent. However, SseM-HA or SseM^V103A^ did not complement the Δ*sseM* mutant strain in the mixed infection (CI= 2.137 ± 0.979, 1.394 ± 0.253, respectively). In contrast, SseM^SEN^ did complement the Δ*sseM* mutant strain in the mixed infection (CI= 0.885 ± 0.215). These results demonstrate that SseM modulates the growth of *Salmonella* during systemic infection, and this phenotype is dependent on the PBM of SseM.

**Fig.4.**
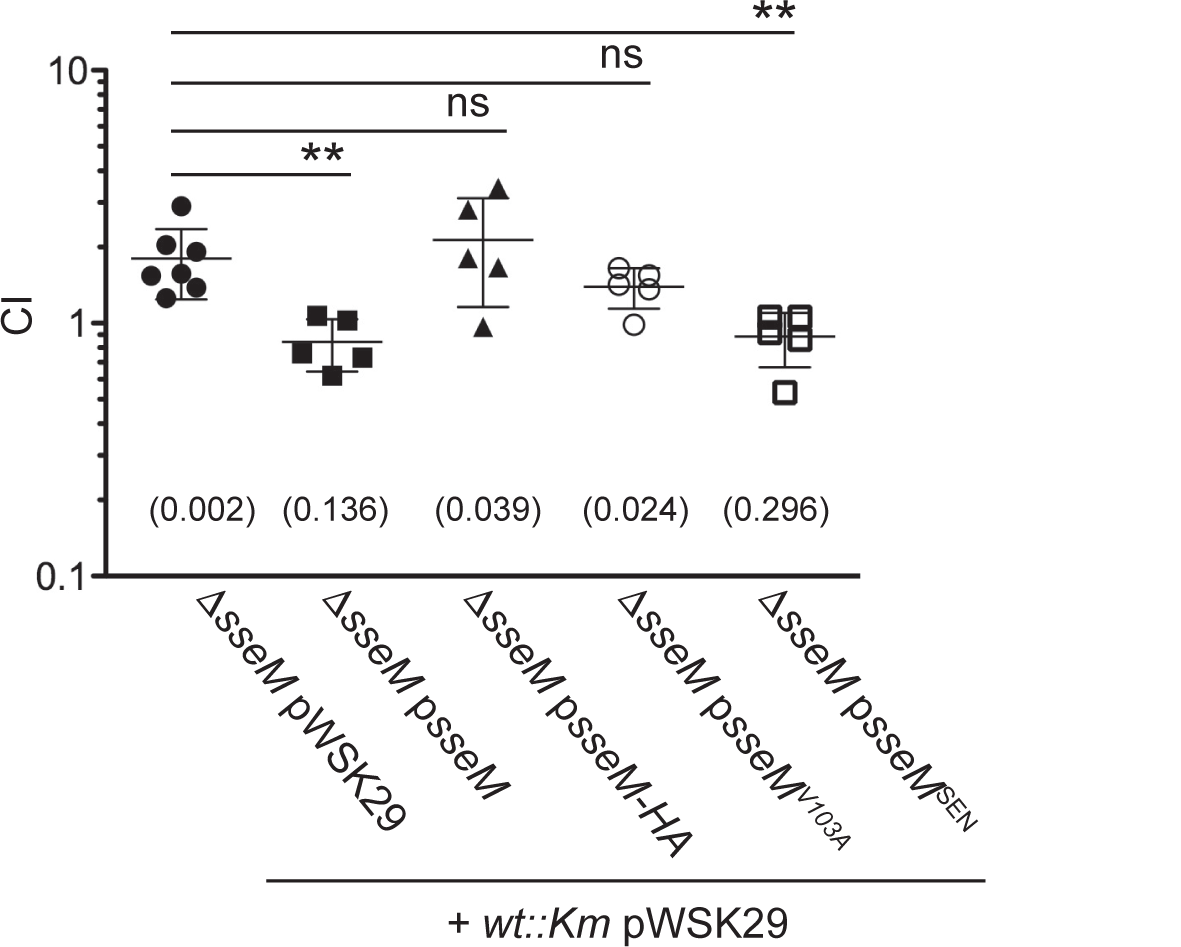
The PBM of SseM is required to downregulate *Salmonella* virulence *in vivo*. BALB/c mice were inoculated by intraperitoneal injection with equal numbers (500 cfu of each of the two strains) of the indicated bacteria. Bacteria were recovered from infected spleens 72 h post-inoculation, and CI values were calculated. The log10 CIs were used for statistical analysis: single sample T-test was used to compare the log10 CI to the hypothetical value of 0 and p value is indicated in the round bracket, one-way ANOVA followed by Dunnett’s multiple comparison test was used to compare with the Δ*sseM* pWSK29/*wt::Km* pWSK29 group (ns: not significant; **: *p* <0.01).

## Discussion

In this work, we investigated the SPI-2 injectisome effector repertoire of the clinical isolate *S.* Enteritidis strain D24359 and identified a previously undescribed effector, which we have named SseM. Like Niemann et al [23], we exploited the property of SPI-2 gatekeeper mutants to hypersecrete effectors into SPI-2-inducing culture medium, which was then collected for mass spectrometry analysis. While Niemann et al [23] passed 500 ml of culture supernatant through a column containing solid-phase extraction resin to prepare samples for mass spectrometry analysis, we only needed to concentrate 50 ml of culture supernatant using a centrifugal filter and fractionated the concentrated samples by SDS-PAGE to prepare samples for mass spectrometry analysis. Our approach therefore provides an easy and cheap method to prepare multiple samples for investigating the SPI-2 effector repertoire from other serovars of *S*. enterica. Eighteen effectors were identified in both our study and that of Niemann et al [23] and a further five unique effectors were found in each study. The genes of certain effectors like *steE* and *sspH1* are not present in D24359. The congruity between these two studies suggests that most, if not all, the SPI-2 repertoire has been identified for these two strains. However, it remains possible that some effectors might be expressed and secreted in low amounts or subject to regulatory control that is absent from *in vitro* growth conditions and still await discovery.

Our previous analysis revealed that all serovars of *S. enterica* subspecies *enterica* have a set of ‘core’ effectors (SseF, SseG, PipB, SteA, SifA, SteD and PipB2) [7]. Here we showed that the newly identified effector SseM is not only present in all serovars of *S. enterica* subspecies *enterica* but is also present in all other four subspecies of *S. enterica* - hence we conclude that SseM is an eighth ‘core’ effector.

Although SseM is annotated as a 110-residue hypothetical protein in most *Salmonella* databases (sFig. 1) we showed experimentally that *sseM* encodes a protein of 103 amino acids, which is translocated by the SPI-2 injectisome and under control of the *pipB2* promoter and the two-component regulatory system SsrAB (Fig. 2B and [20]). This is supported by RNAseq data which only revealed transcription start sites before the *pipB2* gene [27], leading us to conclude that SseM and PipB2 are encoded in the same operon.

SseM, when expressed in isolation in human cells or after translocation by intracellular *Salmonella*, interacted specifically with components of DAPC signalosome [29, 30]: β-2-syntrophin, utrophin/ dystrophin, α-catulin, α-dystrobrevin and β-dystrobrevin. Of particular interest, we identified a PDZ binding motif within SseM and found that both this and the PDZ domain of β-2-syntrophin were required to mediate the interaction between SseM and components of the DAPC signalosome. Both DAPC and DLG1 are involved in several key cellular functions that include not only cell signalling from the adrenergic receptor but also regulation of the cell’s cortical cytoskeleton, cell migration and formation of focal adhesions [30, 35, 36] as well both DLG1 and DAPC regulating tight junctions of polarised epithelial cells [37, 38]. We hypothesise that via its PBM, SseM interferes with one or more of these processes. To our knowledge, SseM is unique among *Salmonella* effectors in containing a functional PBM and as a bacterial protein targeting the DAPC. Several viral oncoproteins target DLG1 to regulate viral virulence [39, 40]. It therefore now essential to investigate the biochemical consequences and physiological significance of SseM’s interaction with DLG1 and DAPC components.

There are several examples of bacterial effectors whose function is mediated through short linear motifs that mediate protein-protein interactions. These include three other PBM-containing effectors (Map, OspE and NleG8) characterised in enteropathogenic *Escherichia coli* [41, 42], *Shigella flexneri* [43], *Citrobacter rodentium* and enterohemorrhagic *Escherichia coli,* respectively [44], with each effector/PBM sequence required for virulence of the corresponding pathogens [41–44]. We found that the Δ*sseM* mutant strain slightly outcompeted the wt strain in the *S.* Typhimurium/ mouse systemic infection model. This modulation of *Salmonella* growth was dependent on the functional PBM of SseM, suggesting that an interaction between SseM and the DAPC acts to restrain bacterial replication during growth in infected tissues. AvrA [45] and SteC [22, 46] are two other effectors whose absence leads to a slight growth advantage of *Salmonella*. The fact that several such mutants exist points to an important aspect of bacterial virulence that remains to be understood.

## Materials and Methods

### Bacterial strains and growth conditions

Bacteria were grown in Luria Bertani (LB) medium supplemented with carbenicillin (50 μg ml^-1^), kanamycin (50 μg ml^-1^) or chloramphenicol (10 μg ml^-1^), for strains resistant to these antibiotics (Ap^r^, Km^r^ and Cm^r^, respectively). To induce SPI-2 gene expression and SPI-2 dependent secretion, bacteria were grown in MgM-MES at pH 5.0 with the corresponding antibiotics when appropriate.

The λ Red recombination system [47] was used to construct the following mutants: *S.* Enteritidis strain D24359 derivatives Δ*spiC::Km* mutant and Δ*spiCssaC::Km* mutant (Primers are listed in Supplemental Table 1), *S.* Typhimurium strain 12023 derivatives Δ*sseM::Km* mutant, Δ*pipB2^promoter^::Km* mutant and Δ*pipB2::Km* mutant. When necessary, pCP20 was used to remove the antibiotic resistance cassette. *S.* Typhimurium strain 12023 derivatives *ssrA::mTn5* mutant [12] and Δ*ssaV::aphT* mutant [2] were described previously.

### Plasmids

Complementing plasmids were constructed by ligating HindIII and PstI-digested plasmid p*ssaGpr* (Ap^r^) [6], a pWSK29 [48] derivative containing the DNA sequence of *ssaG* promoter, with the corresponding digested PCR products: p*sseM*, p*sseM^ACG^*, p*sseM-HA* and p*sseM^V103A^*by using *S.* Typhimurium 12023 genomic DNA as PCR template, and p*sseM^SEN^*by using *S.* Enteritidis D24359 genomic DNA as PCR template.

PciI and NotI-digested M6pblast-GFP (Ap^r^) [49] was ligated with NcoI and NotI-digested PCR products (Supplemental Table 1 for primers and corresponding gene) to construct GFP-tagged effector transfection plasmids: using *S.* Typhimurium 12023 genomic DNA as PCR template for making p*gfp::spvD*, p*gfp::sseM*, p*gfp::sseM-HA*, p*gfp::sseM^RE2A^*, p*gfp::sseM^T101A^*and p*gfp::sseM^V103A^*; using *S.* Enteritidis D24359 genomic DNA as PCR template to make p*gfp: :sseM^SEN^*.

PCR products of *sseM* or *spvD* replaced *steD* gene of pCG36 (Km^r^) to make p*flag::sseM* and p*flag::spvD*, and replaced *steD* gene of pCG189 (Ap^r^) to make p*mCherry::sseM* and p*mCherry::sseM^V103A^*, respectively.

A codon-modified form of SNTB2 gene, eliminating an internal NotI digestion site was synthesised by Invitrogen, and subcloned to PciI and NotI-digested M6pblast-GFP to make plasmid p*gfp::SNTB2*. Overlapping PCR was carried out to amplify SNTB2^GLGI4A^, SNTB2^H176Y^ and SNTB2^H176V^; PciI and NotI-digested PCR products were cloned to plasmid M6pblast-GFP to make p*gfp::SNTB2^GLGI4A^*, p*gfp::SNTB2^H176Y^*and p*gfp::SNTB2^H176V^*.

All the plasmids constructed in this study were verified by DNA sequencing.

### Preparation of secreted proteins for mass spectrometry analysis and immunoblotting

Bacteria were grown overnight in 5 ml LB broth. 1 ml culture was pelleted, washed once with MgM-MES at pH 5.0, and subcultured into 50 ml of MgM-MES at pH 5.0 prior to 6h incubation at 37°C, 200 rpm. Bacteria were pelleted at 10,000 × *g* for 10 min at 4°C, the supernatant was filtered through a ϕ0.2 μm membrane (Acrodisc Syringer Filter, 0.2 μm Supor Membrane, Low protein binding, non-pyrogenic, PALL Life Science) followed by concentration to approximately 200 μl on an Amicon^®^ Ultra-15 Centrifugal Filter with Ultracel-3k membrane (UFC9003, Millipore) at 4°C. 50 μl of concentrated supernatant was run approximately 1 cm into a 12% SDS-PAGE separating gel. The 1cm gel slice, stained with PageBlue Protein Staining Solution (Thermo Fisher Scientific) was sent for mass spectrometry analysis at the Institute of Biochemistry and Biophysics (IBB) at the Polish Academy of Sciences, Warsaw, Poland. Acquired spectra were compared to our annotated *S.* Enteritidis D24359 sequence using the MASCOT search engine.

For pH shift analysis, the subculture was grown for 4 h at pH 5.0 and switched to MgM-MES at pH 7.2 for another 1.5 h. The whole bacterial lysate and secreted fraction were prepared as described previously [5] to make 10 µl of whole bacterial lysate equal to 0.1 OD_600_ of culture and 10 µl of secreted fraction equal to 0.6 OD_600_ of culture. Antibodies used in this study are listed in Supplemental Table 2.

### Bioinformatic analysis

Sequencing data for *S*. Enteritidis strain D24359 has been published previously (ENA accession: ERR037572); however, no genome assembly or annotation was published. To this end, we have downloaded the reads and evaluated their quality using FastQC v0.11.6. The reads were determined to be quality- and adapter-trimmed. Following this, short read assembly was performed using Unicycler v0.4.5. The resulting assembly had 668 contigs and N50 of 10,609. To improve the annotation, we have applied Ragout v2.0 with 4 reference-quality Enteritidis genomes (A1636: GCF_015241115.1, CP255: GCF_015240995.1, D7795: GCF_015240855.1, and P125109: GCA_015240635.1). This resulted in a much more contiguous assembly (2 contings, N50 4,705,460) with 200 kb (∼5%) of the assembly represented as N’s because of the ambiguity in the syntenic blocks.

The resulting assembly was annotated using Prokka v1.12 against a custom *Salmonella* protein database that contained 234,913 unique *Salmonella* proteins annotated using RefSeq Identical Protein Groups. The produced annotation contained 4,448 putative protein-coding genes. The predicted proteins were used as a reference during the mass spectrometry analysis. The code and files necessary to reproduce the assembly and annotation are available at the repository https://github.com/apredeus/D24359.

Protein BLAST was used to search the presence of D24359_01053 in *S. bongori*, *S. enterica* subspecies *salamae*, *arizonae, houtenae*, *indica* and several common serovars of *S. enterica* subspecies *enterica*. ‘Identical Proteins’ in other *S. enterica* serovars were identified and the protein sequences were aligned with Clustal Omega (https://www.ebi.ac.uk/Tools/msa/clustalo/).

To compare the different SseM protein sequence types among *Salmonella* serovars, the complete *Salmonella* genomes were downloaded from Enterobase by searching “Complete Genome” in the “Status” field, which represents the highest assembly quality with circular chromosomes and plasmids (https://enterobase.warwick.ac.uk/, accessed on 2023/06/30). The SISTR1 result from Enterobase were used to identify the subspecies and serovars of the genomes. Only 834 genomes that belong to *Salmonella enterica* subspecies I were included in the analysis.

The *sseM* (*stm2779*) nucleotide sequence from *Salmonella* Typhimurium LT2 (RefSeq: GCF_000006945.2) was used as a reference. A BLAST database was constructed from the *sseM* sequence. Each of the 834 *Salmonella* genomes was queried against the *sseM* database using BLASTn v2.14.0+ [50]. The aligned DNA sequences were then extracted and translated into protein sequences using Seqkit v2.4.0 [51]. The unique SseM protein sequences were summarized and aligned using Clustalo v1.2.4 [52].

To visualise the SseM types in conjunction with the genomic diversity of the *Salmonella* genomes, an MStree of the 834 complete *Salmonella* genomes was generated on Enterobase using the cgMLST scheme with the MSTree2 algorithm [53]. The tree was visualised with Grapetree [26]. In the cgMLST scheme, clusters of genomes with fewer than 900 allele differences are uniform for serovars [53]. Therefore, in Grapetree, the nodes with fewer than 900 allele differences are collapsed into bubbles to visualize the serovars.

AlphaFold Colab Multimer [54] was used to predicate the complex between SseM and the PDZ domain (R^114^ – E^199^) of β-2-syntrophin.

### Bacterial infection, translocation analysis and co-immunoprecipitation

HeLa cells and HEK293 cells were maintained in Dulbecco’s modified Eagle’s medium (DMEM) supplemented with 10% foetal bovine serum (Sigma) at 37°C in 5% (V/V) CO_2_. Infection with *S.* Typhimurium was done as described previously [15].

For translocation assays, HeLa cells were infected for 5 h, then proteasome inhibitor MG132 (Sigma) was added to a final concentration of 10 μg/ml and cells incubated for another 3 h. For immunoblotting analysis, cells were collected, washed once with cold PBS and lysed for 15 min on ice with 50 μl of 0.1%Triton X-100 in PBS. Soluble fraction (containing translocated effectors) was separated from insoluble fraction (containing bacteria and nucleus) by centrifugation at 16,000 × *g* for 10 min at 4°C. Alternatively, cells on glass coverslips were infected as above, fixed, immuno-labelled and analysed with a Zeiss 710 confocal microscope as described [15].

To immuno-precipitate SseM, HeLa cells seeded in a ϕ15 cm petri dish were infected for 14.5 h, then MG132 was added to a final concentration of 10 μg/ml and cells incubated for another 3 h. After a PBS wash, cells were resuspended into 800 μl of buffer A (50mM Tris pH 7.5, 150mM NaCl, 0.5% sodium deoxycholate, 1% Triton X-100, 1mM EDTA and cOmplete, Mini, EDTA-free Protease Inhibitor Cocktail (Roche)) and lysed for 30 min on ice. The lysate was centrifuged at 16,000 × *g* for 10 min at 4°C, supernatant was incubated with 40 μl of Protein G Agarose (Pierce) on a roller at 4°C. The pre-cleaned supernatant was then incubated with 30 μl of Protein G Agarose pre-bound with 50 μl of rabbit anti-SseM antibody for 3.5 h on the roller at 4°C. The agarose was washed 4 times with buffer A and resuspended into 30 μl of 2 × protein loading buffer.

### Generation of stable cell lines and SNTB2 knockout cell lines

Lipofectamine^TM^ 2000 (Invitrogen) was used to transfect HeLa cells with plasmid. Transfected cells were selected with blasticidin to establish stable HeLa cell lines expressing GFP or GFP::SseM.

Two different guide RNAs targeting the coding sequence 871^st^ −891^st^ nt (GGTGTGGATAGCTACGAACC) or 1072^nd^ −1092^nd^ nt (TGCTCTATGACTGTATGCCG) of SNTB2 were cloned onto pX330 [55] with annealed oligos XJY361/362 or XJY363/364 to construct plasmids p361 [KO1] or p363 [KO2] respectively. 24 h after transfection, HEK293 cells were seeded into 96-well plates at 0.3 cells per well. Single clones were screened by immunoblotting with mouse anti-Syntrophin antibody.

### Immunoprecipitation from stable cell lines or transfected cells

GFP-trap agarose (Chromotek) or anti-Flag M2 affinity gel (Sigma) were used to immuno-precipitate GFP-tagged protein or Flag-tagged protein by using buffer B (5% glycerol, 0.5% Triton X-100, 1 mM phenylmethylsulfonyl fluoride (PMSF), PBS) as lysis buffer and buffer C (5% glycerol, 0.1% Triton X-100, 1 mM PMSF, PBS) as washing buffer.

One ϕ10 cm dish of HeLa stable cell line expressing GFP or GFP::SseM, or one ϕ10 cm dish of HEK293 cells transiently transfected with 3 μg plasmid DNA p*flag::sseM* or p*flag::spvD* for 16 h were used for immunoprecipitation. After 4 washes with buffer C, the beads were washed twice with PBS before sending for mass spectrometry analysis. Acquired spectra were compared to database of *Homo sapiens* (Uniprot) using the MASCOT search engine.

For immunoblotting analysis, HEK293 cells seeded in one well of 6-well plate was transfected with 1.5 μg plasmid DNA or co-transfected with 0.75 μg of each plasmid for 16 h before collecting cells for GFP-trap immunoprecipitation. After 4 washes with buffer C, the beads were then resuspended into 30 ml of 2 × protein loading buffer.

### Mouse ethics statement

Animal experiments were performed in accordance with ASPA and UK Home Office regulations. The project licence for animal research (P2ED1F62A) was approved by Imperial College London Animal Welfare and Ethical Review Body (ICL AWERB). Prior to experimentation, mice were given at least one week to acclimatise to the ICL Animal Research Facility.

### Mouse mixed infection

The virulence of *S.* Typhimurium strain 12023 derivative wt*::Km* strain is indistinguishable from wild-type *S.* Typhimurium strain 12023 (J. Poh and D. W. Holden, unpublished), and was used as wild -type strain for competitive index (CI) studies. Female BALB/c mice (7-8 weeks) mice were inoculated intraperitoneally with a mixture of two strains comprising 500 colony-forming units of each strain in PBS, and the CIs were determined from spleen homogenates 72h post-inoculation as described previously [34].

Single sample T-test was used to compare the log10 CI to the hypothetical value of 0 (the value of 0 means that two strains grew equally well *in vivo*). One-Way ANOVA corrected by Dunnett’s multiple comparison test was used to compare the log10 CI to that of the Δ*sseM* pWSK29/ wt*::Km* pWSK29 group.

### Data and materials availability

The code and files necessary to reproduce the assembly and annotation of D24359 are found here https://github.com/apredeus/D24359. All other data are available in the main text of supplementary materials. Correspondence and requests for materials should be addressed to Xiu-Jun Yu: and Teresa L.M Thurston.

## Acknowledgments

A Biotechnology and Biological Sciences Research Council David Phillips Fellowship BB/R011834/1 and ERC starting grant funded by the Engineering and Physical Sciences Research Council EP/X02377X/1 to TLMT and a Wellcome Trust Investigator Award, 209411/Z/17/Z to DWH funded this work.

We thank members from the Holden, Thurston and Hinton laboratories for feedback, and Dr. Camilla Godlee for providing plasmids pCG36 and pCG189.

**sFig.1 Alignment of SseM from different subspecies and different serovars of *Salmonella enterica*.** (A) Alignment of D24359_01053 and STM2779. Identities: 95/103=92%; positives: 97/103=94%. (B) Alignment of SseM protein sequences. Accession numbers for non-*enterica* subspecies are highlighted in grey. EAW1721535.1: *S. enterica* subsp. *indica*; WP_080161159.1: *S. enterica* subsp. *arizonae*; WP_072157437.1: *S. enterica* subsp. *salamae*; WP_071651510.1: *S. enterica* subsp. *houtenae*; WP_011233152.1: serovar Paratyphi A; WP_077905756.1: serovars Agona, Indiana, Goldcoast, Brancaster, Senftenberg; WP_023166795.1: serovars Kentucky, Senftenberg and Gallinarum; WP_077909235.1: serovars Muenchen, Litchfield, Manhattan; WP_077910820.1: serovars Paratyphi B, Java; WP_077463764.1: serovars Heidelberg, Newport, Infantis, Hadar etc. EDQ7330352.1: serovar Paratyphi C; HCK6744786.1: serovars Anatum, Eko, Typhi; WP_014343883.1: serovars Typhimurium, Saintpaul, Paratyphi B, Berta, Bareilly; WP_077945811.1: serovars Brunei, Newport; WP_016701746.1: serovar Enteritidis.

**sFig.2 Alignment of SseM variants**

Sequence alignment of SseM variants from Fig. 1D.

**sFig.3 Alignment of *stm2779*, *sseM_2_pseudo* and *sseM_6_pseudo*.** The extra ‘C’ (underlined bold font) in *sseM_2_pseudo* results in a stop codon (red fonts) after the predicated 30^th^ amino acid. The mutation of the predicated 84^th^ codon TGG to TAG (red font) in *sseM_6_pseudo* results in a truncated protein missing the last 28 residues of SseM.

**sFig.4 Alignment of *stm2779* and *D24359_01053*.** Green fonts indicate the predicated start codon (TTG) and actual start codon (ATG) of *stm2779*. Plasmid p*sseM^ACG^*was constructed by changing the ATG (green fonts) to ACG (red fonts) on p*sseM*.

**sFig.5 Predicted secondary structural features of β-2-syntrophin PDZ domain with AlphFold Colab Multimer.** The secondary structure of PDZ domain (italic fonts) are indicated. Mutated residues of PDZ domain of β-2-syntrophin used in Fig. 3F are highlighted in green.

